# Targeting mitochondrial metabolism with CPI-613 in chemoresistant ovarian tumors

**DOI:** 10.1101/2024.05.16.594585

**Authors:** Mary P Udumula, Faraz Rashid, Harshit Singh, Tim Pardee, Sanjeev Luther, Tanya Bhardwaj, Km Anjaly, Sofia Piloni, Miriana Hijaz, Radhika Gogoi, Philip A Philip, Adnan R Munkarah, Shailendra Giri, Ramandeep Rattan

## Abstract

**Background:** There is evidence indicating that chemoresistance in tumor cells is mediated by the reconfiguration of the tricarboxylic acid cycle, leading to heightened mitochondrial activity and oxidative phosphorylation (OXPHOS). Previously, we have shown that ovarian cancer cells that are resistant to chemotherapy display increased OXPHOS, mitochondrial function, and metabolic flexibility. To exploit this weakness in chemoresistant ovarian cancer cells, we examined the effectiveness of the mitochondrial inhibitor CPI-613 in treating preclinical ovarian cancer.

**Methods:** Chemosensitive OVCAR3, and chemoresistant CAOV3 and F2 ovarian cancer cells lines and their xenografts in nude mice were used. Functional metabolic studies were performed using Seahorse instrument. Metabolite quantification was performed using LC/MS/MS.

**Results:** Mice treated with CPI-613 exhibited a notable increase in overall survival and a reduction in tumor development and burden in OVCAR3, F2, and CAOV3 xenografts. CPI-613 suppressed the activity of pyruvate dehydrogenase and alpha-ketoglutarate dehydrogenase complex, which are two of its targets. This led to a reduction in OXPHOS and tricarboxylic acid cycle activity in all 3 xenografts. The addition of CPI-613 enhanced the responsiveness of chemotherapy in the chemoresistant F2 and CAOV3 tumors, resulting in a notable improvement in survival rates and a reduction in tumor size as compared to using chemotherapy alone. CPI-613 reduced the chemotherapy-induced OXPHOS in chemoresistant tumors. The study revealed that the mechanism by which CPI-613 inhibits tumor growth is through mitochondrial collapse. This is evidenced by an increase in superoxide production within the mitochondria, a decrease in ATP generation, and the release of cytochrome C, which triggers mitochondria-induced apoptosis. Our study demonstrates the translational potential of CPI-613 against chemoresistant ovarian tumors.

## Background

Ovarian cancer ranks as the eighth most common cancer among women globally and is the fifth leading cause of cancer-related fatalities in women [1]. The treatment for ovarian cancer, specifically high grade serous ovarian cancer, includes major cytoreductive surgery followed by standard of care chemotherapy consisting of carboplatin and paclitaxel or neoadjuvant chemotherapy followed by cytoreduction and additional chemotherapy, based upon the patient status [2]. Approximately 75% of patients with advanced stage disease will recur, either with refractory chemoresistant tumors or tumors that will eventually develop chemoresistance [2, 3]. Chemoresistance is a significant challenge in the treatment of ovarian cancer, necessitating the development of innovative medicines to effectively counteract the aggressive growth of tumors [3–5].

Reprogramming of the metabolism is now a well-established hallmark of cancers [6], where dysregulated cellular energetics is a key feature of metabolic rewiring [7, 8]. Increasing data shows that adapting to aerobic glycolysis in cancer cells does not remove the reliance on oxidative phosphorylation (OXPHOS) [9, 10]. OXPHOS has been demonstrated to be the primary energy source for various cancers, with active mitochondria playing a crucial role in several tumor characteristics, including tumor growth, metastasis, recurrence, response and resistance to drugs and preservation of cancer stem cells (CSCs) [11–16]. We have previously demonstrated that ovarian cancer cell lines that are resistant to chemotherapy exhibit heightened metabolic activity, displaying increased OXPHOS through an augmented tricarboxylic acid (TCA) cycle, when compared to cell lines that are sensitive to chemotherapy [16]. We have also shown that when sensitive cells are exposed to chemotherapy, they shift from using glycolysis to employing OXPHOS, and this change is associated with their sensitivity [16]. These studies provide evidence of the connection between chemoresistance and mitochondrial function. A promising new approach to therapy are metabolic inhibitors, particularly those that target mitochondria [17].

We chose the mitochondrial inhibitor CPI-613 (devimistat) to investigate if inhibiting mitochondrial function can alleviate chemoresistance in ovarian cancer models. CPI-613 is a lipoic acid analog that blocks 2 crucial points in the TCA cycle: the pyruvate dehydrogenase complex (PDHc), which facilitates the entry of pyruvate, and the alpha-ketoglutarate dehydrogenase complex (αKGDC), which facilitates the incorporation of glutamine into the TCA cycle [18–21]. Preclinical studies have shown the anti-tumor effect of CPI-613 in lung [22], ovarian [20], pancreatic [21, 23, 24], head and neck [25], clear cell sarcoma [26] and colorectal cancers [27], attributing the mechanism to be inhibition of both PDH and αKGDH, culminating in the suppression of the TCA cycle and mitochondria. CPI-613 has undergone rigorous evaluation in many clinical trials to assess its safety and effectiveness. Although the initial trials of CPI-613 in combination with chemotherapy showed promise [19, 28–30], subsequent larger trials in pancreatic cancer and acute myeloid leukemia failed to deliver the anticipated survival benefits [31–33]. Nevertheless, subsequent analysis of clinical trials demonstrated a statistically significant and beneficial rise in a specific group of patients when the modified FOLFIRINOX regimen was combined with the treatment for pancreatic cancer and in older patients acute myeloid leukemia [24, 33]. Therefore, CPI-613 retains promise, provided that its specific suitability for certain populations and types of tumors has been determined.

In this study, we assess and illustrate the effectiveness of CPI-613 in preclinical models of ovarian cancer that are sensitive and resistant to chemotherapy. Our findings indicate that CPI-613 induces apoptosis in tumor cells via mitochondrial collapse. The research provides compelling data for assessing the effectiveness of CPI-613 in ovarian cancer patients, namely those who have demonstrated resistance to chemotherapy.

## Methods

### Cell lines and reagents

OVCAR3 is a human adenocarcinoma cell line used as a chemosensitive cell line and CAOV3, F2 human epithelial ovarian cancer cells were used as a resistant model for the study. CAOV3 and OVCAR3 cells were purchased from American Type Culture Collection (ATCC, Manassas, VA), cultured in Dulbecco’s modified Eagles media (Hyclone), 5% fetal bovine serum (BioAbChem, Ladson, SC), 4%Insulin-Transferrin-Selenium (Gibco, USA)), 100 U/ml penicillin and 100 U/ml streptomycin (Hyclone), 2 mM L-glutamine, and 1 mM sodium pyruvate. F2 cells transduced to express mCherry (mCherry-F2) [34, 35] were developed and kindly provided by Dr. Gil Mor (Wayne State University, Detroit, MI). These cells were cultured in Roswell Park Memorial Institute base medium, 10% fetal bovine serum, 1% penicillin-streptomycin, 1% HEPES, 1% minimum essential media with non-essential amino acids NEAA and 1% sodium pyruvate. All cells were cultured at 37°C in a humidified 5% CO_2_ incubator. CPI-613 (Devimistat) was obtained from Cornerstone Pharmaceuticals (Cranbury, NJ).

### Animal model and experiments

#### Induction of intra-peritoneal ovarian cancer

Female nude mice of 6-8 weeks were acquired from Jackson Laboratory (Bar Harbor, ME) and were maintained under standard optimized conditions in the animal facility at Henry Ford Hospital (Detroit, MI). Prior to the start of the study, mice were acclimated in house for 1 week. To induce ovarian cancer, mice were injected intraperitoneally (IP) with 5 x 10^6^ OVCAR3, F2 and CAOV3 cells in 200 ul of phosphate-buffered saline. Body weight and tumor growth were monitored once a week till the end of the study as previously reported [36]. Mice were sacrificed when the abdominal circumference reached 8 cm or when they exhibited partial impaired mobility.

#### Induction of subcutaneous ovarian cancer

Female nude mice of 6-7 weeks age obtained from Jackson Laboratory were acclimatized for 1-2 weeks. To induce subcutaneous tumors, 2 x 10^6^ cells/200 µl F2 cells were injected subcutaneously into the left flank of each mouse. When tumors grew an average volume of 100 mm^3^, mice were randomly grouped into control (vehicle corn oil) or CPI-613, chemotherapy and chemotherapy and CPI-613 combination. Tumor volume was calculated as described previously [37].

#### Treatments

All treatments were started after 1 week of tumor injection. All CPI-613 mice were injected with 2.5 mg/kg body weight in corn oil for 4 weeks post-tumor injections for 3 days in a week. For chemotherapy mice received 20 mg/kg body weight of carboplatin and 50 mg/kg of paclitaxel for 6 cycles, 2 doses a week for 3 weeks.

#### Survival curve

For estimating overall survival, 7 or 9 mice per group were injected with respective cells and tumors were allowed to proceed until the abdominal circumference reached 8 cm for IP tumors. For the subcutaneous model, mice were euthanized when the tumors reached a size of 1000 mm^3^, according to the Henry Ford Health Institutional Animal Care and Use Committee approved end point. The mice were then humanely euthanized. Survival curves were generated using Kaplan-Meir analysis using Prism 10 (GraphPad Software, La Jolla, CA).

#### In situ fluorescence imaging of tumors

The mCherry fluorescence signals of the tumor were recorded using the Xenogen IVIS system 2000 series (Perkin Elmer, Akron, OH). Images were acquired over a period of 20 minutes with a 15-second exposure. Living Image software (Perkin Elmer) was used to read and integrate the total fluorescence signals in each region. Fluorescence measurements were quantified as total photon flux emission as photons/second in the images acquired at the same exposure time for the various groups [38].

### Ethics statement

All protocols were approved by the Henry Ford Hospital Institutional Animal Care and Use Committee prior to any experiments (#1472). All institutional and national guidelines for the care and use of laboratory animals were followed.

### Alamar blue viability assay

CAOV3, F2 and OVCAR3 cells (2000 cells/well) were cultured in the presence and absence of various doses of CPI-613 ranging from 5 to 250 µm. After 72 hours of incubation at 37°C and 5% CO_2_, Alamar blue was added to the cultures according to the manufacturer’s protocol (BioRad, Hercules, CA). After incubation for 4 hours, absorbance was measured using a SYNERGY H1 Hybrid reader. For estimating growth of various fuels, cells were plated in Dulbecco’s Modified Eagle Media containing high pyruvate (1 mM) or low pyruvate (0.1 mM) and high (2.5 mM) or low glutamine (0.5 mM). Alamar blue assay was performed at 0-72 hours of respective media exposure. The test was run in triplicate and repeated thrice. The results were expressed as percentage of cell viability.

### Seahorse metabolic analysis

OVCAR3, CaOV3 and F2 cells or single cell suspension of tumor cells isolated from CaoV3 and F2 xenografts were plated at a density of 7 x10^4^ cells/well in cell-Tak coated XFe 96 cell plates. Oxygen consumption rate (OCR) and extracellular acidification (ECAR) rate were measured using XFe 96 seahorse bioanalyzer (Agilent Seahorse XF Analyzers, Santa Clara, CA) and analyzed as describe before [16, 36, 38, 39].

### Real-time polymerase chain reaction

RNA extraction was performed as described previously [39–41]. In brief, RNA was extracted from cultured cells or tumors of control and CPI-613 treated mice using an RNA assay kit (Qiagen, Germantown, MD) and quantified by Qubit Fluorimeter (Thermo Fisher Scientific, Waltham, MA). The cDNA was synthesized using 1 µg of total RNA using high quantity cDNA kit in 20 μl reaction mixture using CFX BioRad Real-time PCR (polymerase chain reaction) Detection System. Ribosomal protein L27 was used as a housekeeping gene. All the primers were purchased from Integrated DNA Technologies (Coralville, IA) and Real time Primers (Melrose Park, PA). All primers used in the study are detailed in the supplementary material (Table S1).

### Western blotting

Western blotting was performed as published before [36, 38]. In brief, after indicated treatments, the total protein was isolated from tumor cells after lysing cells in a protein lysis buffer (50 mM Tris-HCl, pH 7.5, 250 mM NaCl, 5 mM EDTA, 50 mM NaF, and 0.5% Nonidet P-40; containing a protease inhibitor cocktail; Sigma-Aldrich, St Louis, MO), and quantified using BCA (Bicinchononic acid assay kit; Thermo Fisher). An equal amount of protein was separated by 4-20% sodium dodecyl sulphate polyacrylamide gel electrophoresis and transferred onto the polyvinylidene difluoride membrane and blocked with 5% skimmed milk. Antibodies used included anti-DLST, PDHa, cytochrome C, apoptotic protease activating factor 1 (APAF1), cleaved caspase 3, cleaved caspase 9, cleaved PARP and β-actin. All antibodies were purchased from Cell Signaling Technologies (Danvers, MA) and were diluted at a ratio of 1:1000 in 5% BSA for primary antibody incubation.

### Immunohistochemistry

Tumor tissue was fixed in 10% formaldehyde for 48 hours and paraffin embedded. Sections were cut and processed for immunohistochemistry to detect Ki-67. The Ki-67 antibody was purchased from Abcam (Cambridge, MA) and used at a concentration of 1:200. Solutions obtained from Dako Cytomation (Carpinteria, CA) were used for performing immunostaining, using the Dako Autostainer Link 48 as described previously [38]. A minimum of 3-6 slides per group and 6-10 fields per slide were evaluated under a high-power field using Nikon eclipse Ci-S microscope (Melville, NY).

### Quantitation of TCA metabolites by LC-MS/MS

The levels of TCA metabolites in tumor cells of untreated, CPI-613, chemotherapy and chemotherapy and CPI-613 combination mice were quantified by ultrahigh performance liquid chromatography-tandem mass spectroscopy (LC-MS/MS) (Waters, Milford, MA) as described before [37, 42]. *Chemicals and reagents.* TCA cycle standard mix 1 and 2 (cat no. MSK-TCA1 and MSK-TCA 2A) and its ^13^C-labeled metabolites were purchased from Cambridge Isotope Laboratories, Inc. (Tewksbury, MA). Acetonitrile (HPLC-grade), formic acid, MS-grade water and methanol were purchased from Sigma-Aldrich.

#### TCA metabolites LC-MS method for extraction and quantification

As per experimental design, all the metabolites were extracted using 80:20 acetonitrile and water mixture. TCA standards were reconstituted in (water and acetonitrile, 20:80) to prepare 10 mg/ml stock solutions. Linear standards curve at the range of 5-1000 ng/ml of each analyte (citric acid, aconitase, iso-citric acid, ±-ketoglutaric acid, itaconate, α-ketoglutarate, succinic acid, fumarate, and malic acid) was produced from stock solutions in the relevant matrix. Each standard (100 µl) was mixed to obtain a 7-point standard calibration curve within a range of 5-1000 ng/ml. Each internal standard (10 µg/ml) (isotope-labelled TCA mixtures standards mix Sets 1 and 2A) were spiked into all samples as an internal standard (final concentration) to allow quantification based on the ratio of the internal standard to each intermediate peak. Approximately, 3 million cells were rapidly rinsed by 2 mL of 37°C deionized water to the cell surface. Cells were collected in 2-ml tubes after washing, followed by adding 500 µL of extraction solution (acetonitrile and water [80:20]) and stored at −80°C for further processing. 10 µL of each isotope-labeled TCA mixtures standards mix (Sets 1 and 2A) was added as an internal standard followed by vortex to mix. Three cycles of extraction were carried out by vertexing for 1 minute followed by sonicating for 1 minute (each). After sonication, samples were kept at −20°C for protein precipitation, followed by centrifugation at 15,000 rpm for 20 minutes at 4°C. Supernatants were loaded into pre-conditioned Phenomenex Strata XL-100 60 mg/3 ml cartridges (Torrance, CA) and passed through it using positive pressure manifold (Agilent Technologies). Flow through was collected subsequently and 100 µL of filtrate was mixed with 100 µL of water for the LC-MS/MS system. The TCA cycle and its intermediate metabolites were identified and quantified based on retention time and m/z match to injections of authentic standards, retention time and accuracy bases and were quantified using internal standard area for the respective metabolites. Phenomenex Luna-NH2, 2.0 x 150 mm, 3 µm column was used to achieve an optimal separation of all TCA intermediates. The flow rate of 0.25 ml/min was used with a mobile phase A (10 mM NH_4_OAc buffer at pH 9.8) and mobile phase B (acetonitrile). Column temperature was optimized at 25°C for the best chromatographic separation. The metabolites were eluted from the column at a flow rate of 0.25 ml/min using a gradient system (mobile phase B): 80% (0.01–0.5 min), 80→20% (0.5–18.0 min), 20% (18.0–19.5 min), 20 → 80% (19.5–22 min), and re-equilibration (22–25 minutes). TCA cycle intermediates were monitored using negative MRM (multi reaction monitoring) polarity. Identification was achieved based on retention time and MS/MS ion conformation. Seven calibration standards ranging from 5-1000 ng/ml were subjected to the full extraction procedure 3 times before analysis. The mean correlation coefficients of each metabolite were linear r2 > 0.99 and were obtained (n = 3) from 5-1000 ng/ml. The calibration curve, prepared in a control matrix, was constructed using peak area ratios of the calibration samples by applying a one/concentration weighting (1/x) linear regression model. All quality control sample concentrations were then calculated from their PARs against the calibration curve. The parameters for triple quadrupole detector mass spectrometry equipped with an electrospray ionization probe: Capillary, voltage, 3.5 kV for negative mode: Source temperature 120°C: Desolvation temp: 450°C; Cone gas flow: 150 L/h: Desolvation gas flow:1000 L/Hr Collision gas flow: 0.25 mL/min and nebulizer gas flow: 7 Bar.

#### Data analysis

Mass spectrometric data was acquired by MassLynx v4.2software (Waters, Milford, MA. TargetLynx software was used for preparing the calibration curve and absolute quantitation of all TCA metabolites in the samples. Analyte concentrations were calculated using a 1/x weighted linear regression analysis of the standard curve.

### Mitochondrial dysfunction detection

#### Mitochondrial membrane potential

Single cell suspension of the xenografts was prepared using Accutase (Innovative Cell Technologies, San Diego, CA). Tumor cells were labeled with Ep-CAM (APC/Cyanine7 anti-mouse CD326 Antibody, BioLegend, San Diego, CA) for identification. MitoProbe TMRM (tetramethylrhodamine methyl ester) kit for flow cytometry (Invitrogen, # M20036) was used as per manufacturer’s instructions to measure the membrane potential. Attune NXT flow cytometer (Thermo Fisher Scientific, Waltham, MA) set to excite and emit light at 488/575 nm was used for measurements.

#### Mitochondrial superoxide detection

Single cell suspension of the xenografts was prepared using Accutase (Innovative Cell Technologies). Tumor cells were labeled with Ep-CAM as above for identification and treated with MitoSOX red dye (Thermo Fisher) for 15 minutes, washed 3 times, and then analyzed using an Attune NXT flow cytometer with excitation and emission wavelengths of about 396/610 nm. For microscopic detection, cells were treated with CPI-613 at a concentration of 100 μM for 48 hours. Subsequently, 5 μM of the fluorogenic dye MitoSOX Red was applied to the cells on coverslips to measure mitochondrial superoxide levels, and the cells were incubated for 15 minutes. The cells were then washed and counterstained with DAPI (4′,6-diamidino-2-phenylindole) for imaging. Intensity of MitoSOX positive cells were measured using Nikon NIS Imaging software (Melville, NY).

#### Measurement of lipid peroxidation

Lipid peroxidation was measured by detection of malondialdehyde in the tumor lysates using Lipid Peroxidation kit (AbCaM, #ab118970) as per the manufacturer’s instructions.

#### Mitochondrial metabolism array

A human mitochondrial energy metabolism PCR array (Qiagen RT2 Profiler PCR Arrays, PAHS-008YA) was performed from cDNA synthesized from 1 µg of total RNA from the tumors of control and CPI-613 treated mice. This library profiles 84 genes involved in mitochondrial energy metabolism. The quantitative expression of the gene was calculated from the cycle threshold value of each sample and was normalized using the housekeeping genes by using online PCR quantification software provided by the manufacturer [43].

### Statistical analysis

An unpaired t-test or one-way analysis of variance was used where appropriate. Kaplan Meier analysis was used to determine the survival curve. The significance of survival curves was estimated by using the Gehan Breslow-Wilcox test and the Mantel-Cox tests. All analyses were carried out using Prism 10 (GraphPad Software, La Jolla, CA).

## Results

### Chemoresistant ovarian cancer cells are dependent on OXPHOS

We have previously shown that chemoresistant ovarian cancer cells are metabolically energetic with increased OXPHOS [16]. To confirm our previous observations, we performed carboplatin sensitivity and bioenergetic profiling of chemosensitive (OVCAR3) and chemoresistant (F2 and CAOV3) cell lines. F2 and CAOV3 exhibited resistance to carboplatin (Fig. 1A), an increased oxygen consumption rate with differential ECAR compared to chemosensitive OVCAR3 (Fig. 1B-E), confirming our previous observations [16] of chemoresistance correlating with increased OXPHOS and a highly metabolically active phenotype (Fig. 1F). Next, we compared the dependency of sensitive and resistant cell lines on pyruvate and glutamine, the two major fuels for maintaining an upregulated TCA cycle [44, 45]. Sensitive OVCAR3 cells showed no effect on growth when glutamine was reduced and when pyruvate was reduced in media, minimal growth inhibition was seen, indicating that if glucose is present as a carbon source, these can survive and grow (Fig. 1G and 1J). In contrast, both resistant cell lines F2 and CaOV3 showed reduced proliferation under low glutamine or pyruvate conditions (Fig. 1H, 1I, 1K, and 1L), despite glucose being present in the media. Interestingly growth was more dependent on glutamine compared to pyruvate in both F2 and CaOV3 cell lines (Fig. 1K and 1L). This indicates that the increased TCA cycle is being sustained both by pyruvate and glutamine, especially glutamine. Further, we assessed the expression of key enzymes PDHc and αKGDC that facilitate the entry of pyruvate and glutamine into the TCA cycle respectively. Both F2 and CaOV3 cells showed increased mRNA expression of DLST (dihydrolipoamide S-succinyltransferase), the E2 transferase subunit of αKGDC [46] and PDH subunits; PDH1a and PDH1b [47] compared to OVCAR3 (Fig. 1M-O). Higher expression of DLST and PDH1a was further validated at the protein level in the resistant cell lines (Fig 1P-R). Thus, chemoresistant metabolically active ovarian cancer cells upregulate the 2 key nodes of TCA cycle to maintain a high OXPHOS via upregulation of TCA cycle.

**Figure 1:**
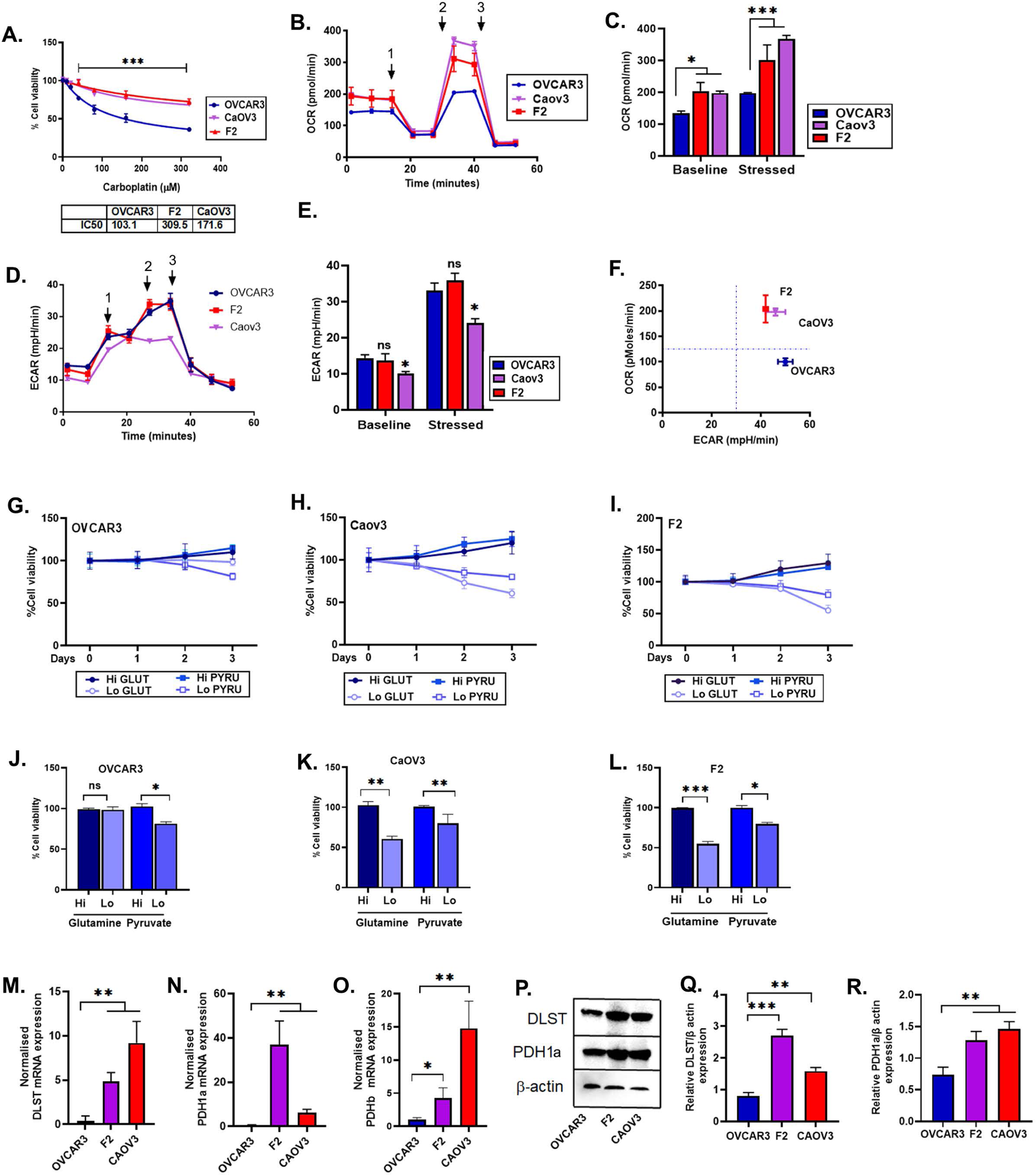
Chemoresistant ovarian cancer cells are dependent on oxidative phosphorylation. (A) OVCAR3, Caov3 and F2 cells (2000 cells/well in triplicates) were plated in 96 well plates and treated with indicated concentrations of CPI-613 and 72 hours later, cell viability was assessed using the Alamar blue viability assay. (B) OVCAR3, Caov3 and F2 cells (70,000 cells/well in triplicates) were subjected to real-time XFe Seahorse analysis for bioenergetics profiling. Oxygen consumption rate (OCR) an indicator for mitochondrial respiration was assessed with port injections of (1) oligomycin, (2) FCCP (Carbonyl cyanide-p-trifluoromethoxyphenylhydrazone), and a combination of (3) rotenone-antimycin. (D) Extracellular acidification rate (ECAR), an indicator of glycolysis, was measured with port injections of (1) glucose, (2) oligomycin, and (3) 2-DG (2-deoxyglucose). (C, E) The bar graph represents basal and stressed OCR and ECAR (n=3). (F) The ratio of basal OCR and ECAR was plotted to ascertain the metabolic phenotype of the cells as per XFe Seahorse analysis. (G, H, I) OVCAR3, CaOV3 and F2 cells (2000 cells/well in triplicates) were plated in 96 well plates and subjected to growth conditions of media containing high pyruvate (1 mM) or low pyruvate (0.1 mM) and high (2.5 mM) or low glutamine (0.5 mM). Cell viability was examined by Alamar blue assay at 0-72 hours of respective media exposure. The test was repeated thrice. (J, K, L) The bar graph represents the final cell viability at 72 hours of various media exposure. (M, N, O) mRNA isolated from all 3 cell lines were subjected to Qpcr to measure the expression levels of (M) DLST; dihydrolipoamide S-succinyltransferase (N) PDH1a; pyruvate dehydrogenase E1 subunit alpha 1 and (O) PDHb; pyruvate dehydrogenase E1 subunit beta. (P) Representative immunoblots showing expression of DLST, PDH1a and beta-actin. (Q, R) Bar plots show normalized densitometry analysis from 2 individual blots.

### CPI-613 inhibits growth and improves overall survival in ovarian cancer xenografts

Because of the increased TCA cycle, OXPHOS and dependency on pyruvate and glutamine, we selected CPI-613, an inhibitor of the TCA cycle that inhibits the 2 key nodes of TCA cycle: PDHc and αKGDC [18, 21, 24]. We evaluated the efficacy of CPI-613 in the sensitive OVCAR3 and resistant F2 and CAOV3 IP xenografts as detailed in the Methods section. Overall survival was significantly improved in all 3 models by CPI-613 as seen by an increase in survival by 11 days in OVCAR3, 12 days in CAOV3 and most profoundly by 18 days in F2 bearing xenograft mice, the most resistant and aggressive cell line (Fig. 2A, 2C, and 2E). The increase in survival was supported by the reduction in tumor weights (Fig. 2B, 2D, and 2F). Histological analysis of the Ki-67 stained tumor tissue indicated CPI-613 significantly reduced the number of proliferating tumor cells compared to vehicle treated xenografts in all 3 models (Fig. 2G-L). Thus, CPI-613 is effective in inhibiting both chemotherapy sensitive and resistant ovarian tumors.

**Figure 2:**
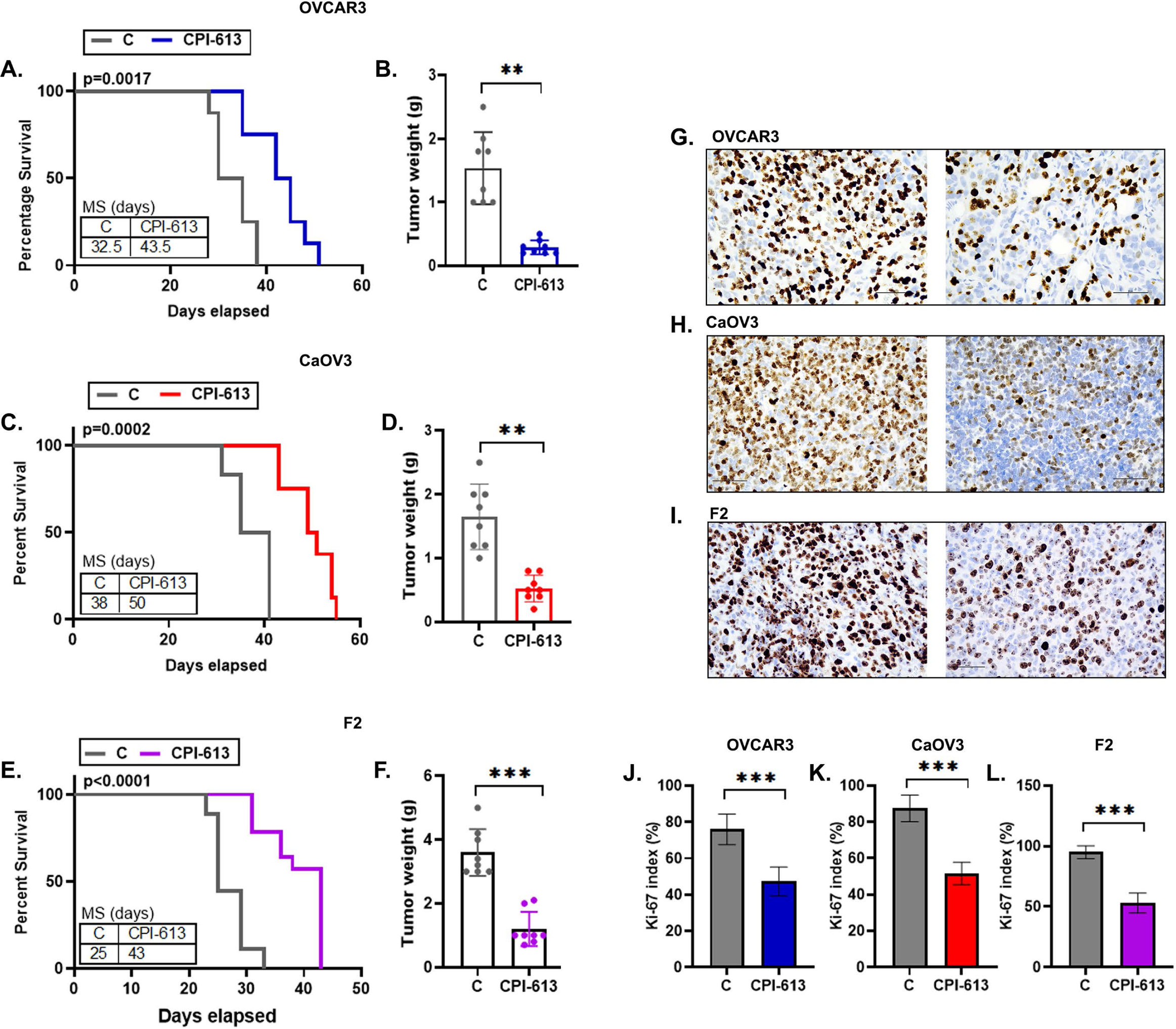
CPI-613 inhibits growth and improves overall survival in ovarian cancer xenografts. (A) OVCAR3, (C) CaOV3 and (E) F2 intraperitoneal xenografts (n=8) were treated with 2.5 mg/kg body weight of CPI-613 thrice a week in corn oil and observed for survival as described in the Methods section. Control groups were treated with corn oil alone. The significance of the Kaplan Meier graph was estimated by Gehan-Breslow-Wilcoxon test. MS = median survival in days. Bar graph shows average tumor weights isolated from the (B) OVCAR3, (D) CaOV3 and (F) F2 mouse models (n=8). Representative immunohistochemistry images (400x) showing Ki67 stain in (G) OVCAR3, (H) CaOV3 and (I) F2 xenografts. (J, K, L) Bar graphs show the Ki-index calculated from *n* = 5 images per group at high-power field (HPF).

### CPI-613 inhibits TCA cycle in ovarian cancer xenografts

To validate that CPI-613 can inhibit the TCA cycle and modulates the energy metabolism of the xenografts, we determined the level of TCA cycle enzymes and mitochondrial activity in CPI-613 treated xenografts. CPI-613 treated tumors from OVCAR3, CaOV3 and F2 xenografts showed a significant decrease in the mRNA expression of its target enzymes; αKGDC subunits: DLST, DLD (dihydrolipoamide dehydrogenase), and OGDH (oxoglutarate dehydrogenase) and both the PDHc subunits: PDH1a and PDH1b (Fig 3A-C). This was also confirmed by decreased protein expression of DLST and PDH1a in the tumors treated with CPI-613 (Fig. 3D-G).

**Figure 3:**
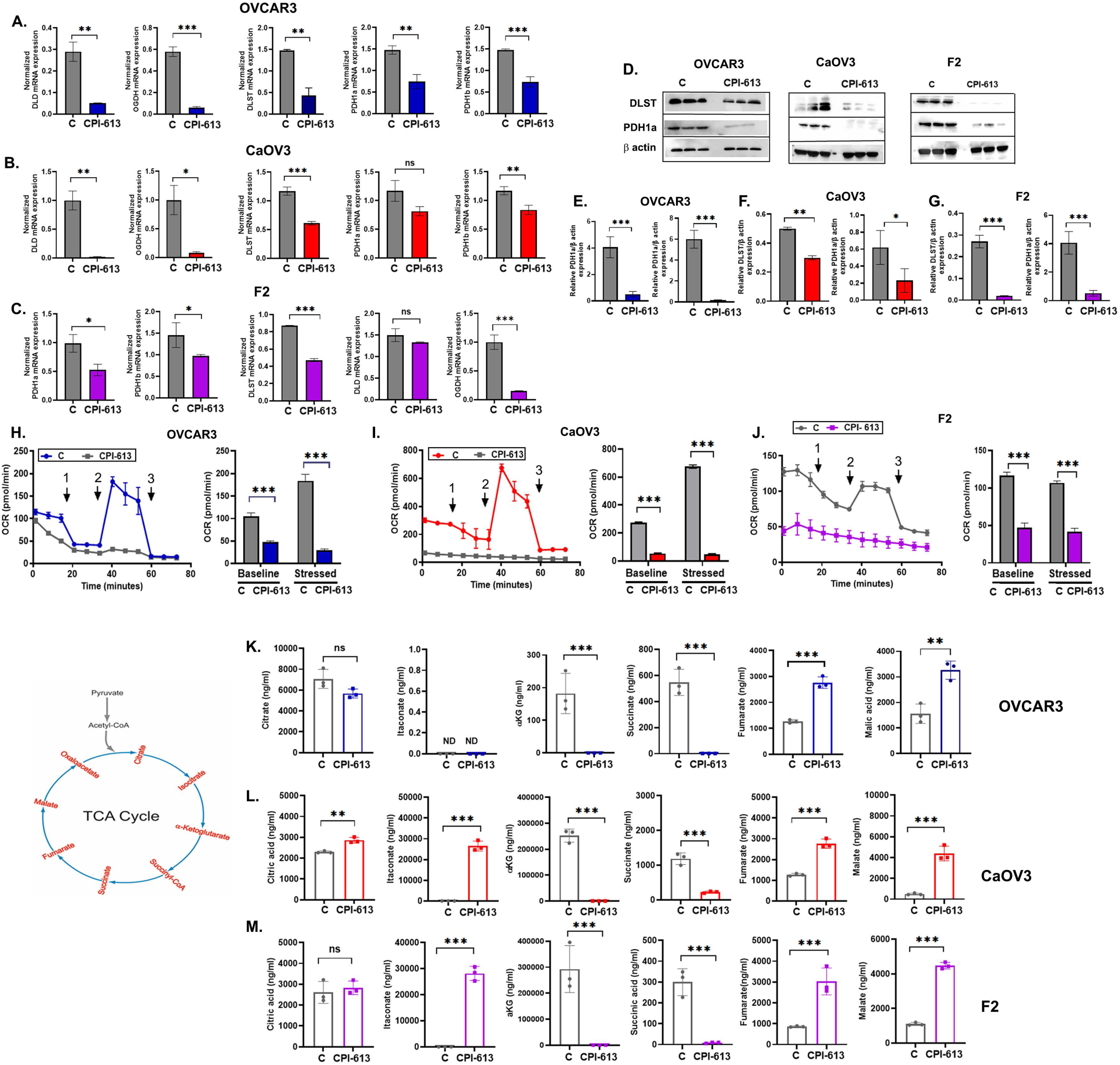
CPI-613 inhibits TCA cycle in ovarian cancer xenografts. Pooled tumor tissue (n=4) was used to isolate mRNA from (A) OVCAR3, (B) CaOV3 and (C) F2 xenografts and subjected to quantitative polymerase chain reaction to estimate the expression levels of dihydrolipoamide dehydrogenase (DLD), oxoglutarate dehydrogenase (OGDH), dihydrolipoamide S-succinyltransferase (DLST), pyruvate dehydrogenase E1 subunit alpha 1 (PDH1a) and pyruvate dehydrogenase E1 subunit beta (PDHb). (D) Representative immunoblots showing expression of DLST, PDH1a and beta-actin in proteins isolated from 3 individual mouse tumor tissue of OVCAR3, CaOV3 and F2 xenografts. (E, F, G). Bar plots show normalized densitometry analysis from 2 individual blots from each cell line. (H-J) Single cell suspension was prepared from freshly isolated xenografts (n=3), cells were pooled to plate 70,000 cells/well in triplicates and subjected to real-time XFe Seahorse analysis for bioenergetics profiling. Oxygen consumption rate (OCR), an indicator for mitochondrial respiration, was assessed with port injections of (1) oligomycin, (2) FCCP (Carbonyl cyanide-p-trifluoromethoxyphenylhydrazone), and a combination of (3) rotenone-antimycin in xenografts from (H) OVCAR3, (I) CaOV3 and (J) F2. The bar graph represents basal and stressed OCR. (K-M) Targeted analysis of the TCA cycle metabolites was performed to assess the levels of various metabolites in pooled xenografts (n=3) in triplicates from (K) OVCAR3, (L) CaOV3 and (M) F2. The schematic figure represents a simplified version of the TCA cycle.

Bioenergetic analysis by Seahorse bioanalyzer of the single cell suspensions of the extracted tumors showed that CPI-613 completely suppressed basal and stressed oxygen consumption rate, compared to vehicle treated tumors, indicating a collapse of energy production by the mitochondria (Fig. 3H-J). No significant change was seen in the ECAR at the baseline or stressed conditions, suggesting that glycolysis was not impacted by CPI-613 (Fig. S1). To further confirm that TCA cycle is impacted by CPI-613, we quantified key TCA metabolites by LC-MS/MS-based analysis [37, 42]. While the levels of key metabolites detected varied between the tumors, there was consistent decrease in levels of αKG (alpha-ketoglutarate) and succinic acid in all 3 tumors, indicating a strong suppression of αKGDC and entry of glutamine into the TCA cycle (Fig. 3K-M). Citrate levels were not altered between control and CPI-613 treated F2 and OVCAR3 tumors, while CAOV3 tumors in the CPI-613 group had increased citric acid levels, suggesting a variable impact on PDHc and pyruvate entry into the TCA cycle (Fig. 3K-M) or presence of an alternate fuel source. Fumarate and malate levels were also significantly increased in CPI-613 treated OVCAR3, CAOV3 and F2 tumors, suggesting synthesis of key metabolites even if the αKG to succinate step is blocked via alternate pathways. Thus, CPI-613 inhibits TCA cycle/mitochondrial respiration and modulates energy metabolism in both sensitive and resistant ovarian tumors.

#### CPI-613 restores chemosensitivity in chemoresistant ovarian xenografts

To determine if CPI-613 can specifically enhance chemotherapy efficacy in the chemoresistant ovarian tumors, we tested the combination of CPI-613 and chemotherapy in the chemoresistant CaOV3 and mCherry-F2 IP xenograft models. Chemotherapy alone improved survival by only 6 days in the CaOV3 xenografts, while a combination of chemotherapy and CPI-613 resulted in a more than 20-day increase with a profound decrease observed in the tumor weights (Fig S2). In mCherry-F2 IP model, chemotherapy alone improved survival by only 4 days in contrast to CPI-613, while the combination with CPI-613, survival was improved by approximately 30 days (Fig. 4A). The survival improvements were reflected in the tumor burden as assessed by final tumor end weights (Fig 4B) and in situ imaging of mCherry in the F2 tumors at 5 weeks post-tumor injections (Fig 4C and 4D). To further test the efficacy of the combination treatment, subcutaneous F2 xenografts were allowed to grow to a volume of approximate 100mm^3^, prior to randomizing and starting treatments. While chemotherapy alone showed minimal abatement in the tumor growth, both CPI-613 alone and CPI-613 and chemotherapy combination groups showed a significant decrease in tumor growth (Fig. 4E) and tumor weight (Fig. 4F). This translated to an enhanced overall survival of the combination group (Fig. 4G), when compared to chemotherapy alone group, as well as CPI-613 alone group. Survival in the combination group was improved by 31 days compared to chemotherapy alone and by 14 days compared to CPI-613 alone (Fig 4G). The impressive inhibition in tumor growth was mirrored in the in-situ imaging of the mCherry in the F2 tumors at 5-weeks post-therapy (Fig 4H and I). Thus, CPI-613 alone is effective against aggressive chemoresistant ovarian tumors and can also enhance the efficacy of chemotherapy against chemoresistant ovarian tumors.

**Figure 4:**
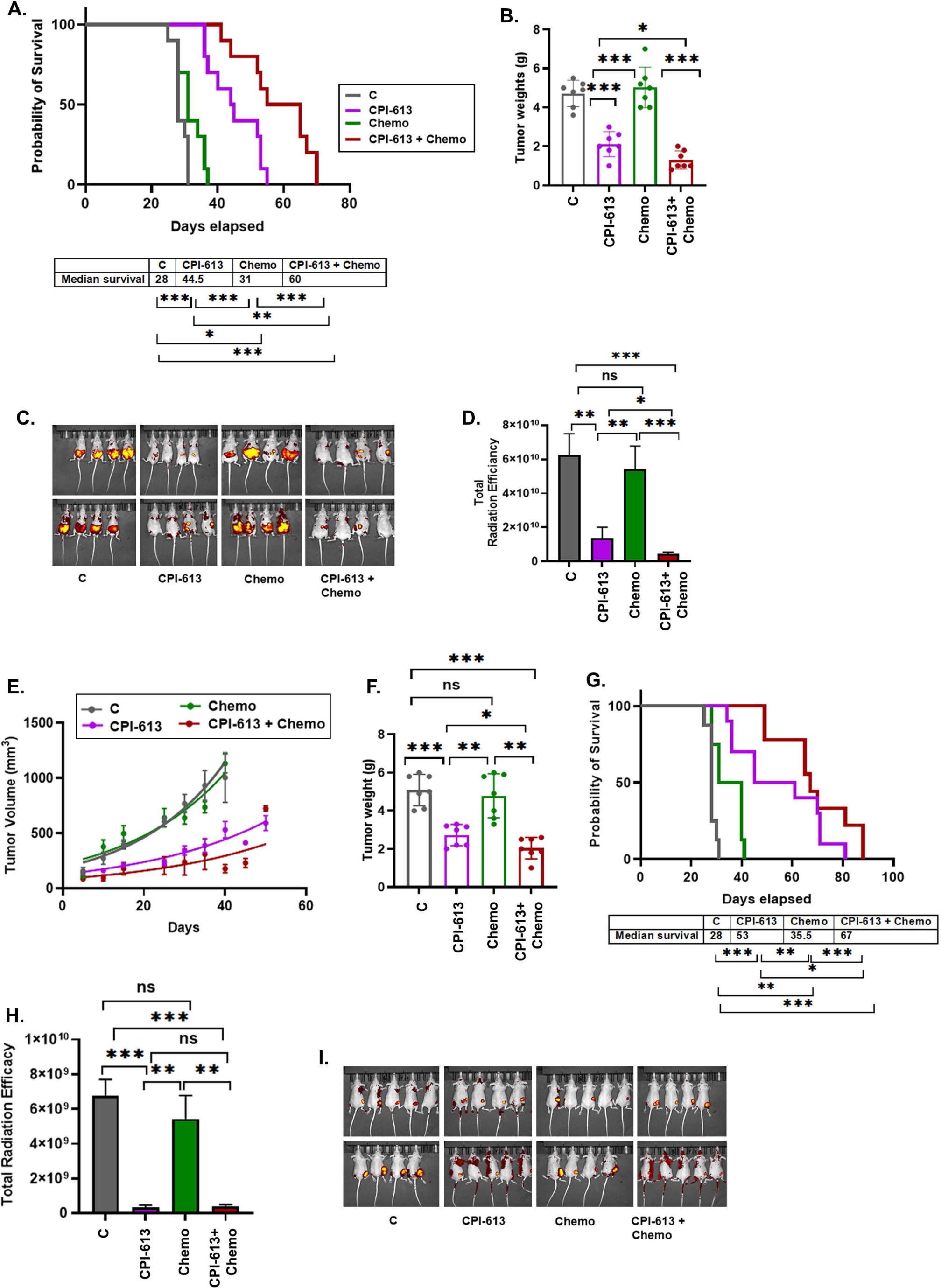
CPI-613 restores chemosensitivity in chemoresistant ovarian xenografts. (A) mCherry-F2 intraperitoneal xenografts (n=8) were treated with CPI-613 alone, chemotherapy (Chemo) alone and a combination of both (CPI-613 + Chemo) and observed for survival as described in the Methods section. The significance of the Kaplan Meier graph was estimated by Gehan-Breslow-Wilcoxon test. (B) Bar graph showing isolated average tumor weights. (C) Representative mCherry bioluminescence pictures indicative of tumor burden. (D) Quantification of the fluorescence signals was done by measuring total photon flux emission/second at the same exposure time for the various groups. mCherry-F2 subcutaneous xenografts (n=7) were allowed to grow to an average size of 100 mm^3^ before treating with CPI-613 alone, chemotherapy (Chemo) alone and a combination of both (CPI-613 + Chemo). (E) Trajectory of tumor growth as represented by measurement of tumor volume. (F) Bar graph showing isolated average tumor weights at the end of the study. (G) Kaplan Meier graph representing the survival of mice. The significance of the Kaplan Meier graph was estimated by Gehan-Breslow-Wilcoxon test. (H) Quantification of the fluorescence signals was done by measuring total photon flux emission/second at the same exposure time for the various groups shown as representative mCherry bioluminescence pictures indicative of tumor burden (I).

### CPI-613 reverses the chemotherapy induced metabolic reprogramming in chemoresistant ovarian xenografts

We and others have previously demonstrated that platinum chemotherapy can induce mitochondrial respiration, a highly active metabolic phenotype and is associated with development of chemotherapy resistance [16, 48, 49]. Seahorse bioanalyzer profiling of the fresh single cell suspension of the xenografts isolated from F2 and CaOV3 treated tumors showed that chemotherapy treated tumors had increased OCR in both models (Fig 5A, B, F, and G), while ECAR was increased in F2 tumors (Fig S3). CPI-613 effectively inhibited the chemotherapy induced bioenergetic increases of both OCR (Fig 5A, B, F, G) and ECAR (Fig S3), indicating its ability to reverse the chemotherapy induced bioenergetic adaptations in ovarian tumors. Measurement of the TCA cycle metabolites showed that chemotherapy did not affect the key downstream metabolites of αKDGC; αKG and succinate in both CaOV3 (Fig 5D and 5E) and F2 (Fig 5I and 5J) tumors. Similarly, acetyl coenzyme A (acetyl-CoA), the downstream metabolite of PDHc, was not impacted by chemotherapy but surprisingly increased by CPI-613 in both tumor models (Fig 5C and 5H), indicating a source other than pyruvate contributing to acetyl-CoA synthesis as a compensatory mechanism [50]. Overall, CPI-613 can inhibit chemotherapy induced mitochondrial respiration and TCA cycle reprograming.

**Figure 5:**
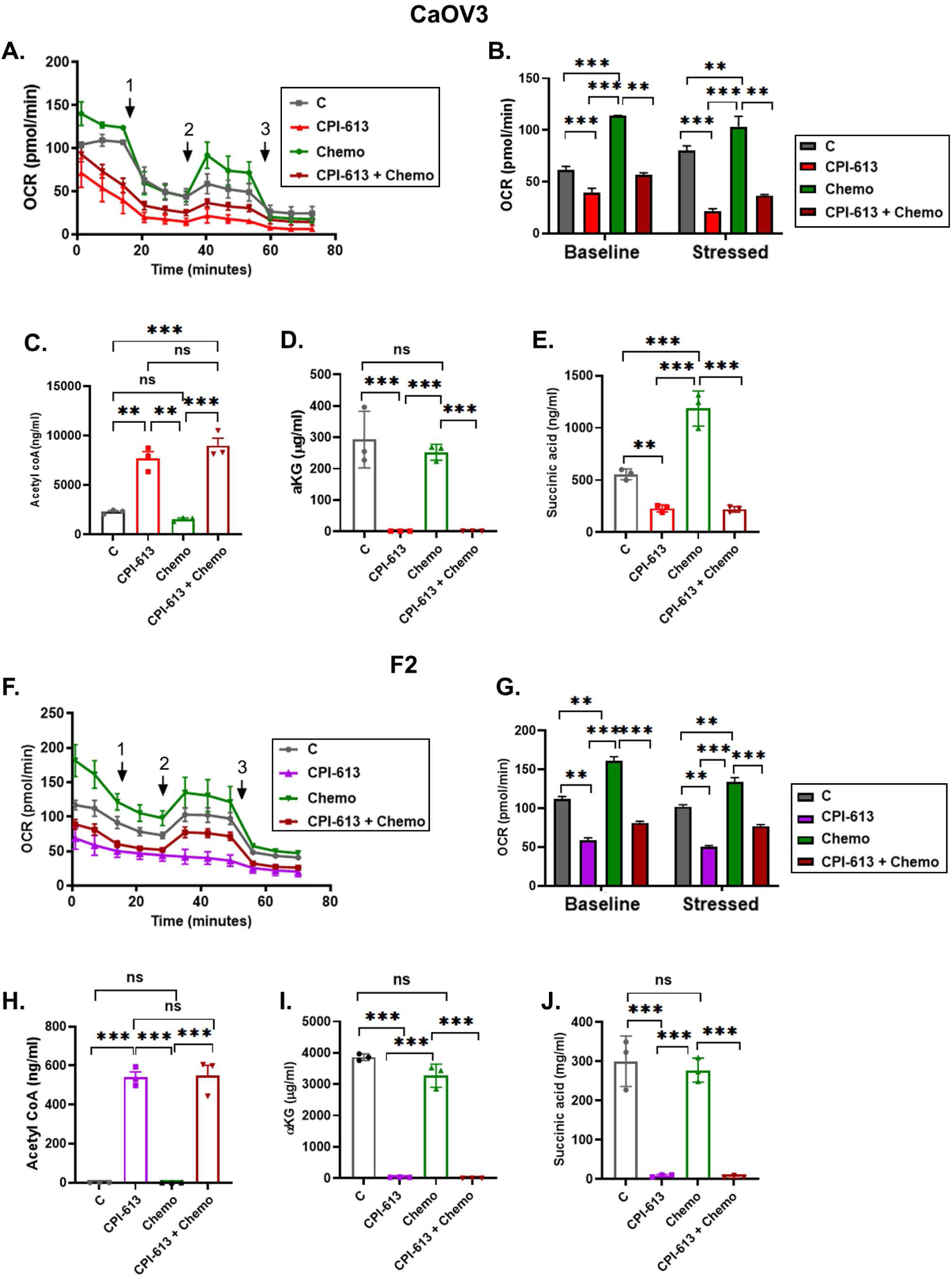
CPI-613 reverses the chemotherapy induced metabolic reprogramming in chemoresistant ovarian xenografts. Single cell suspension was prepared from freshly isolated (A) CaOV3 and (F) F2 xenografts (n=3), cells were pooled to plate 70,000 cells/well in triplicates and subjected to real-time XFe Seahorse analysis for bioenergetics profiling. Oxygen consumption rate (OCR), an indicator for mitochondrial respiration, was assessed with port injections of (1) oligomycin, (2) FCCP, and a combination of (3) rotenone-antimycin. (B, G) The bar graph represents basal and stressed OCR. Targeted analysis of the TCA cycle metabolites was performed to assess the levels of various metabolites in pooled xenografts (n=3) in triplicates. Key metabolites are shown as (C, H) acetyl CoA, (D, I) alpha-ketoglutarate (αKG), and (E, J) succinate.

### CPI-613 causes mitochondrial collapse in chemoresistant ovarian xenografts

Since we observed a drastic decrease in OXPHOS in response to CPI-613, we postulated that CPI-613 may be causing a complete mitochondrial collapse. Single cell suspension from xenografts were measured for mitochondrial dysfunction. We first measured the mitochondrial membrane potential as it represents the electrochemical gradient between the interior and exterior of the mitochondria and is a sign of healthy mitochondria and cells [16, 51]. Ep-CAM positive tumor cells from CPI-613 treated CaOV3 and F2 xenografts showed significant decrease in TMRE florescence indicating mitochondrial membrane depolarization (Fig. 6A-C and 6E-G), which was further supported by decreased ATP levels (Fig 6D and 6H). Another marker of mitochondrial dysfunction is the generation of superoxide, a toxic product that results in accumulated oxidative damage indicated by lipid peroxides or modified DNA bases [52, 53]. Using MitoSox red mitochondrial superoxide indicator, we observed an increase in the fluorescent dye in Ep-CAM positive tumor cells from CPI-613 treated CaOV3 and F2 xenografts, indicating the presence of increased oxidation products (Fig. 6I-L). This was further supported by an increased lipid peroxidation as seen by increased levels of malondialdehyde (MDA)[53], in both xenografts (Fig. 6M and 6N). Microscopic analysis of the ovarian cancer cells stained by MitoSox also validated the increased fluorescence and presence of superoxide in response to CPI-613 (Fig 6O-R). Using mRNA isolated from CPI-613 treated and untreated xenografts, we performed a mitochondrial metabolism PCR array that measured 84 genes crucial for mitochondria metabolism and respiration. We observed that in CPI-613 treated xenografts, majority of the gene expression was downregulated compared to control CaOV3 and F2 tumors (Fig S4). Interestingly, while many similar mitochondrial genes were inhibited in both the xenograft models, the extent of gene expression inhibition between the 2 xenografts was differed (Fig S4).

**Figure 6:**
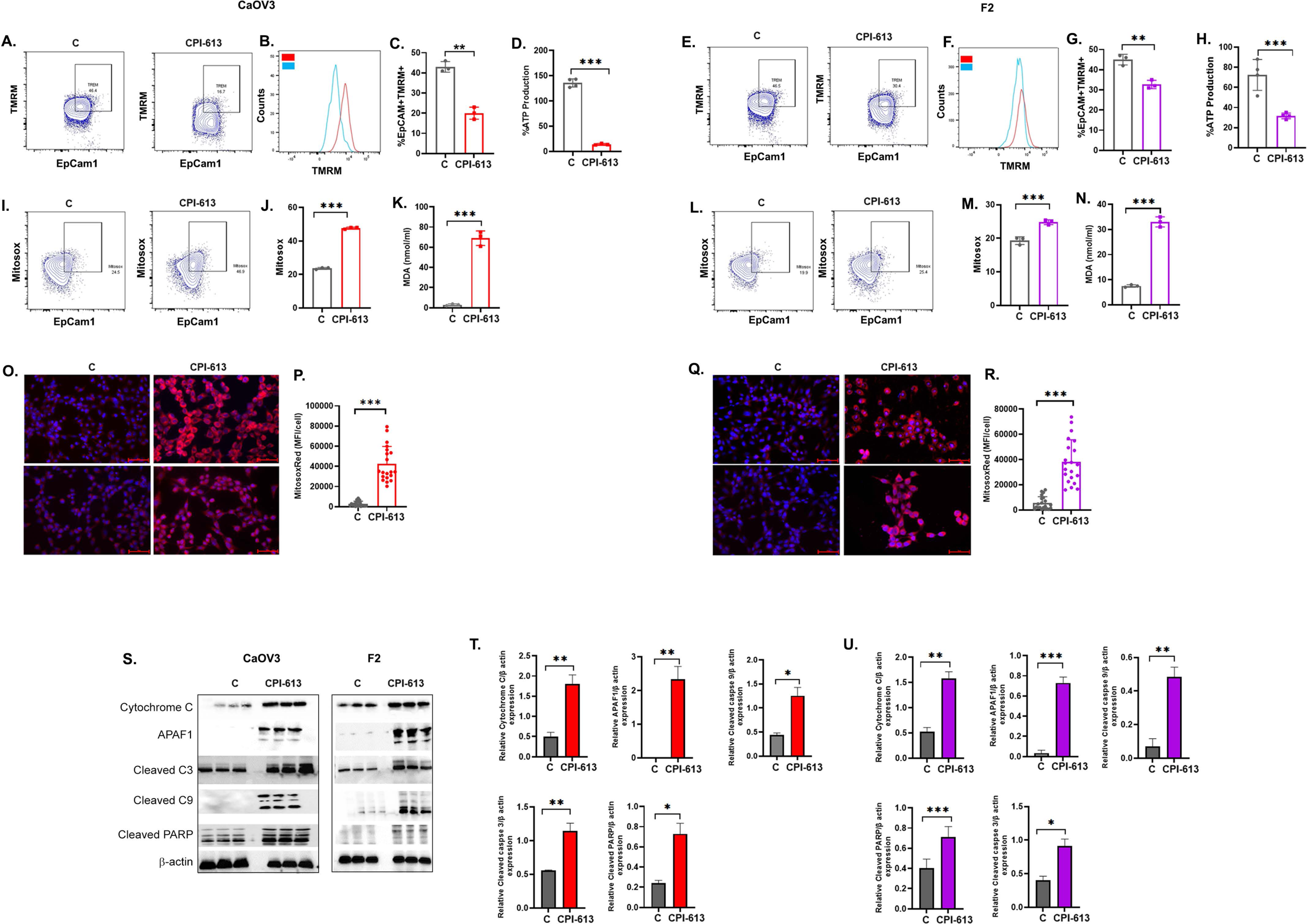
CPI-613 causes mitochondrial collapse in chemoresistant ovarian xenografts. (A & E) Single cell suspensions were prepared from freshly isolated xenografts (n=3), cells were pooled and cells in triplicates were labeled with epithelial cell marker Ep-CAM antibody and Mitoprobe TMRM (tetramethylrhodamine methyl ester) and mitochondria membrane potential was measured using flow cytometry. Representative flow cytometry pictures of Ep-CAM positive cells displaying TMRM fluorescence in (A) CaOV3 and (E) F2 xenografts. (B, F) Representative histograms showing the shift of polarization in cells from both xenografts. (C, G) Quantification of the Ep-CAM positive cells with TMRM measured by flow cytometry. ATP levels were measured as per manufacturer’s instructions in single cell suspensions from (D) CaOV3 and (H) F2 xenografts. Mitosox Red was used to measure the mitochondria superoxide in single cell suspensions from (I, J) CaOV3 and (K, L) F2 xenografts in Ep-CAM positive cells by flow cytometry. Bar graphs represent the quantification of the Mitosox Red signal. Lipid peroxidation was measured by measuring the malondialdehyde (MDA) in single cell suspensions from (M) CaOV3 and (N) F2 xenografts by ELISA. Caov3 cells (O) and (Q) F2 cells were plated on coverslips and treated with CPI-613 (100 μM) for 48 hours and then stained with Mitosox Red and counterstained with DAPI. Representative fluorescence images are shown (400x magnification). (P, R) Bar graphs show quantification of the fluorescence intensity measured in 400x objective field from 3 slides per treatment. (S) Representative immunoblots showing expression of cytochrome C, APAF1 (apoptotic protease activating factor-1), cleaved caspase 3 (cleaved C3), cleaved caspase 9 (cleaved C9), cleaved PARP and beta-actin in proteins isolated from 3 individual mouse tumor tissue of CaOV3 and F2 xenografts. (E, F, G) Bar plots show normalized densitometry analysis from 2 individual blots from each cell lines. (T, U) Bar plots show normalized densitometry analysis from 2 individual blots from each xenograft.

Mitochondrial damage can induce apoptosis by release of pro-apoptotic proteins into the cytoplasm, resulting in mitochondrial mediated apoptosis [54]. CPI-613 treated xenografts showed increased apoptosome proteins [55]: cytochrome C, APAF1 and caspase 9 (Fig. 6S-U), in contrast to untreated control xenografts. The apoptosome formation in response to CPI-613 resulted in final activation of caspase 3 and cell death as observed by an increase in expression of cleaved caspase 3 and PARP (Fig. 6S-U), indicating apoptosis in both F2 and CAOV3 tumors. Thus CPI-613 has the potential to induce mitochondria collapse and the ensuing mitochondrial mediated apoptosis in chemoresistant ovarian tumors.

## Discussion

Ever since metabolic reprogramming was identified as a characteristic of cancer, the importance of mitochondrial metabolism in driving tumor development, tumor metastasis, chemotherapy resistance, and radiation resistance has taken center stage [56]. Previous studies, including our own, have shown that abnormal mitochondria, specifically an overactive TCA cycle, play a significant role in providing metabolic advantages to ovarian cancer cells [14–16]. This metabolic advantage promotes aggressive growth, helps the cells evade therapy, and maintains their CSC-like behavior. These findings suggest that targeting mitochondrial function could be a potential strategy for inhibiting ovarian cancer.

In our current study, we provide evidence of the substantial effectiveness of CPI-613, a lipoate analog that inhibits both PDHc and αKGDC of the TCA cycle, in combating chemoresistant ovarian cancer xenografts. Our initial findings confirmed our previous observation that the chemoresistant ovarian cancer cell lines CaOV3 and F2 are metabolically active, utilizing both glycolysis and higher OXPHOS [16]. This metabolic activity is associated with their reliance on TCA fuels, specifically pyruvate and glutamine, even in the presence of glucose. Additionally, these cell lines showed increased expression of PDHc and αKGDC subunits, in contrast to the chemosensitive OVCAR3 cell line. CPI-613 effectively suppressed the growth of OVCAR3, CaOV3, and F2 xenografts, leading to enhanced overall survival. Notably, it had the most pronounced impact on the F2 cell line, which is characterized by higher aggressiveness and resistance to chemotherapy. Significantly, CPI-613 enhanced the effectiveness of chemotherapy in both the xenografts that were resistant to treatment. The data indicate that CPI-613 has potential as an anti-tumor agent, particularly against chemoresistant ovarian cancer. A recent study showcased the efficacy of CPI-613 in specifically targeting CSCs derived from the ovarian cancer cell lines and xenografts [20]. It also exhibited the potential of CPI-613 to limit the enrichment of CSCs caused by chemotherapy and PARPi. CSCs in ovarian and other malignancies are responsible for the development of recurrent and chemoresistant disease [34, 48]. Multiple studies have demonstrated that CSCs rely on metabolic flexibility for their survival and prefer OXPHOS [34, 48]. A recent publication, which has not undergone peer review, provides evidence of the efficacy of CPI-613 in specifically targeting pancreatic cancer CSCs[57]. Therefore, our finding that the CSC enriched F2 cell is highly affected by CPI-613 aligns with these reports. Nevertheless, we observe a suppression of tumor growth in the xenografts, irrespective of CSCs, leading to an enhancement in survival rates.

Preclinical studies in several types of tumors have verified that the primary mechanism of CPI-613 is the inhibition of PDHc and αKGDC, resulting in the inhibition of the TCA cycle, which affects mitochondrial activity and leads to the death of tumor cells. CPI-613 has the ability to alter lipid metabolism in pancreatic cancer, which triggers AMPK-ACC signaling [23]. This signaling pathway ultimately leads to apoptosis and autophagy [23]. In colorectal cancer, CPI-613 causes cell cycle arrest and apoptosis mediated by Bcl2-like protein 11, Bim [27]. Both investigations also indicated a reduction in mitochondrial activity and the production of reactive oxygen species [23]. Our study also confirms the inhibition of PDHc and αKGDC leading to TCA cycle inhibition and decreased mitochondrial respiration.

Interesting observations were seen when TCA metabolites were measured in xenografts in response to CPI-613. The most consistent was the almost absence of αKG and succinate, indicating a significant impact of CPI-613 on αKGDC inhibition, which would block conversion of αKG to succinyl-CoA leading to succinate [50]. The decrease in αKG suggests that CPI-613 is also able to block glutamine-glutamate-αKG analplerosis [58] into the TCA cycle, but we did not observe any change in the levels of glutamate in the tumor cell (Fig. S5). Alternately, it may indicate an increase in reductive carboxylation of αKG to citrate [58], as citrate levels were unchanged in OVCAR3 and F2 but increased in CaOV3 tumors. The increase in this metabolic route is supported by the increase seen in other metabolites like itaconate from isocitrate [56] and an unexpected increase in acetyl-CoA (Fig. S5) in CaOV3 and F2 tumors, which was predicted to decreased in response to PDHc inhibition by CPI-613. An increase in acetyl CoA may also reflect the compensatory increase in use of fatty acid reserves [50, 58] by the cancer cells to maintain the TCA cycle for survival. While itaconate is a well-known anti-inflammatory and immunomodulator metabolite [25], its role in tumor biology is controversial [59, 60]. On the other hand, acetyl-CoA has been shown to mainly promote tumor growth due to its multiple functions as fuel and transcriptional regulation by chromatin remodeling [61]. Another interesting metabolite change observed was an increase in fumarate, primarily considered an oncometabolite [62], despite the block in succinate synthesis. These data indicate that the intricate overlapping metabolic routes come into play on inhibition of the TCA cycle by CPI-613, perhaps as an attempt to maintain the energy metabolism and survival of the cancer cells. These observations also exhibit the diverse response of each cell type/xenograft in response to the metabolic inhibitor CPI-613, underlining the existence of heterogeneity.

Nevertheless, the significant reduction in OXPHOS and the severe decrease of key TCA cycle metabolites observed in both chemoresistant ovarian tumor cells and those treated with chemotherapy and/or CPI-613, indicates a substantial decline in mitochondrial activity. We verified this by identifying reduction in mitochondrial membrane potential, accompanied by a fall in ATP levels. Additionally, we saw an increase in mitochondrial superoxide production, which correlated with lipid peroxidation. Furthermore, there was a decrease in the expression of almost all genes associated with mitochondrial energy and metabolism. This indicates a total breakdown of mitochondria, which resulted in apoptosis. Our observations confirm the induction of the mitochondrial mediated apoptotic cascade, as evidenced by the increase in apoptosome levels and the activation of caspase 3, leading to PARP cleavage.

## Conclusions

Our work shows that CPI-613 has a significant effect on the TCA cycle and mitochondrial function, ultimately resulting in the death of tumor cells. CPI-613 exhibits a discerning effect on chemoresistant cancer cells, likely because these cells and CSCs rely more heavily on OXPHOS. Although the recent performance of CPI-613 in clinical trials has been disappointing, recent post hoc analysis of trials indicate that select patient populations may be able to benefit from CPI-613. A post-hoc analysis of the acute myeloid leukemia clinical trial indicates a trend of dose response in older patients in contrast to younger patients [33]. This is pertinent to ovarian cancer as it primarily effects elderly women and for whom age constitutes a substantial risk factor [1, 63]. Therefore, CPI-613 exhibits considerable promise as a pharmaceutical candidate deserving of additional investigations, whether utilized in isolation or in conjunction with conventional chemotherapy or targeted therapies, particularly in the case of chemoresistant ovarian cancer.

## List of Abbreviations

αKDGC: alpha-ketoglutarate dehydrogenase complex
αKG: alpha-ketoglutarate
ACC: acetyl-CoA carboxylase
AMPK: AMP-activated protein kinase
APAF1: apoptotic protease activating factor 1
ATP: adenosine triphosphate
DNA: deoxyribonucleic Acid
BSA: bovine serum albumin
CoA: Coenzyme A
CSC: cancer stem cells
CD: cluster of differentiation
DAPI: 4′,6-diamidino-2-phenylindole
DLD: dihydrolipoamide dehydrogenase
DLST: dihydrolipoamide S-succinyltransferase
ECAR: extracellular acidification
EDTA: ethylenediamine tetraacetic acid
Ep-CAM: epithelial cell adhesion molecule
HCl: hydrochloric acid
LC/MS/MS: liquid chromatography tandem mass spectrometry
MDA: malondialdehyde
NaCl: sodium chloride
NaF: sodium fluoride
OCR: oxygen consumption rate
OGDH: oxoglutarate dehydrogenase
OXPHOS: oxidative phosphorylation
PARP: poly-ADP ribose polymerase
PARPi: poly-ADP ribose polymerase inhibitors
PDHc: pyruvate dehydrogenase complex
TCA: tricarboxylic acid
TMRM: tetramethylrhodamine methyl ester

## Declarations

### Availability of data and materials

Not applicable. No datasets were generated during this work.

### Authors’ contributions

M.P.U., H.S., T. B., K.A. and S.P., performed research, analyzed the data, and edited the manuscript; F.R. performed research and analyzed the data; T.P., S. L., M.H., R.G., P.P., A.M. and S.G. participated in data discussions and designing the study and edited the manuscript; R.R. designed and supervised research, analyzed the data and edited the manuscript.

## Acknowledgements

We would like to thank Cornerstone Pharmaceuticals for providing us CPI-613. We would like to thank Ms. Stephanie Stebens for editing and formatting the manuscript.

