## Supplemental File for "Targeting mitochondrial metabolism with CPI-613 in chemoresistant ovarian tumors"

**Table 1: List of primers used.**

| List of Primers | Primer Sequence |
|---|---|
| DLST | Real time primers |
| PDH 1A | F GGA TGG TGA ACA GCA ATC TTG CC<br>R TCG CTG GAG TAG ATG TGG TAG C |
| PDH 1B | F TGT AAC TGT GGA AGG AGG CTG G<br>R CAT CAG CAC CAG TGA CAC GAA C |
| SDH A | F GAG ATG TGG TGT CTC GGT CCA T<br>R GCT GTC TCT GAA ATG CCA GGC A |
| SDH B | F GCA GTC CAT AGA AGA GCG TGA G<br>R TGT CTC CGT TCC ACC AGT AGC T |
| DLD | F CTG AGT GAA GGA GAC TTG CTG G<br>R TCC CAG AGG AAC ATC CCT TGT G |
| OGDH | F GAGGCTGTCATGTACGTGTGCA<br>R TACATGAGCGGCTGCGTGAACA |
| ATP 6 | F GCGCCACCCTAGCAATATCA<br>R GCTTGGATTAAGGCGACAGC |
| DLOOP2 | F CCCTTCCCCATTTGGTCT<br>R TGGTTTCACGGAGGATGG |
| ND4 | F GCTCCATCTGCCTACGACAA<br>R GCTTCAGGGGGTTTGGATGA |
| 16S | F CACTGCCTGCCCAGTGA<br>R ATACCGCGGCCGTTAAA |
| L27 | Real time primers |

**Supplementary legends**

**Figure S1: CPI-613 has modest impact on ECAR in ovarian cancer xenografts.** Single cell suspension was prepared from freshly isolated xenografts (n=3), cells were pooled to plate 70,000 cells/well in triplicates and subjected to real-time XFe Seahorse analysis for bioenergetics profiling. Extracellular acidification rate (ECAR) an indicator of glycolysis was measured with port injections of (1) glucose, (2) oligomycin, and (3) 2-DG in (A) OVCAR3, (C) CaOV3, and (E) F2 xenografts. The bar graph represents basal and stressed ECAR (B, D, F).

**Figure S2: CPI-613 restores chemosensitivity in chemoresistant ovarian xenografts.** (A) CaOV3 intraperitoneal xenografts (n=8) were treated with CPI-613 alone, chemotherapy alone and a combination of both and observed for survival as described in the Methods section. The significance of the Kaplan Meier graph was estimated by Gehan-Breslow-Wilcoxon test. (B) Bar graph showing isolated average tumor weights.

**Figure S3: CPI-613 reverses the chemotherapy induced metabolic reprogramming in chemoresistant ovarian xenografts.** Single cell suspension was prepared from freshly isolated xenografts (n=3), cells were pooled to plate 70,000 cells/well in triplicates and subjected to real-time XFe Seahorse analysis for bioenergetics profiling. Extracellular acidification rate (ECAR), an indicator of glycolysis, was measured with port injections of (1) glucose, (2) oligomycin, and (3) 2-DG in (A) CaOV3 and (C) F2 xenografts. The bar graph represents basal and stressed ECAR (B, D).

**Figure S4: CPI-613 inhibits mitochondrial energy and metabolism gene expression.** Pooled tumor tissue (n=4) was used to isolate mRNA isolated from (A) CaOV3 and (B) F2 xenografts and subjected to mitochondrial energy and metabolism gene array according to manufacturer's instructions. The expression of 84 genes was normalized and analyzed using the software provided by the manufacturer. The data is represented as a dot blot graph showing the fold change in genes. Some of the highly altered gene expression was truncated to fit the graph.

**Figure S5: CPI-613 modulates TCA cycle metabolites in ovarian cancer xenografts.** Targeted analysis of the TCA cycle metabolites was performed to assess the levels of various metabolites in pooled xenografts (n=3) in triplicates from OVCAR3, CaOV3 and F2. Metabolites measured included glutamate (A-C) and acetyl-CoA (D-F).

**Figure S1: CPI-613 has moderate effect on ECAR in ovarian cancer xenografts**

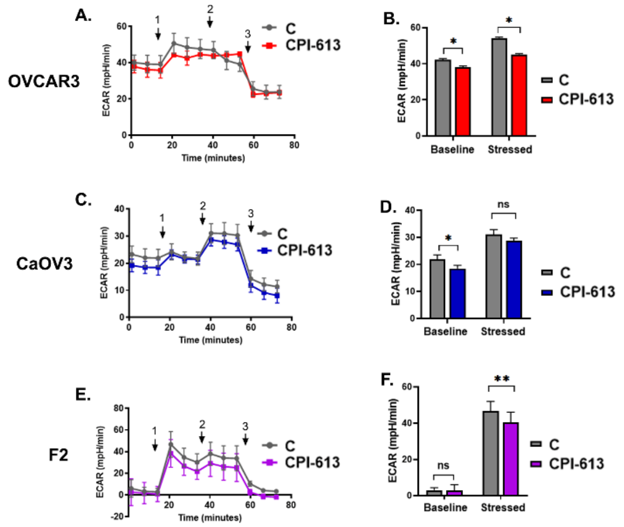

**Figure S2: CPI-613 restores chemosensitivity in chemoresistant CaOV3 xenografts.**

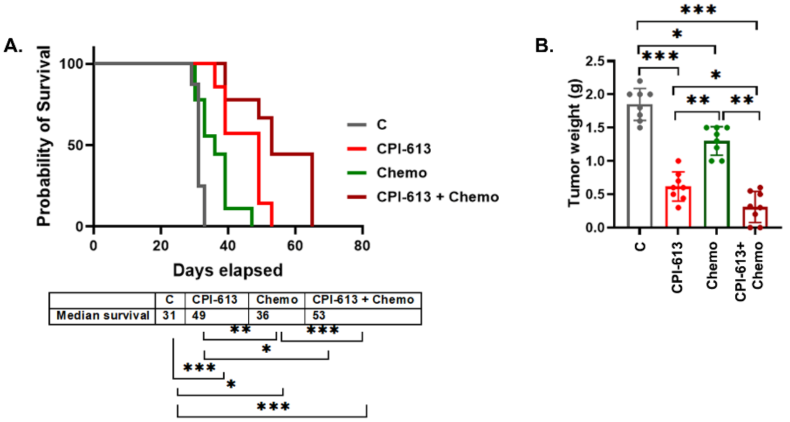


| | C | CPI-613 | Chemo | CPI-613 + Chemo |
|---|---|---|---|---|
| Median survival | 31 | 49 | 36 | 53 |

**Figure S3: CPI-613 reverses the chemotherapy induced metabolic reprogramming in ovarian xenografts.**

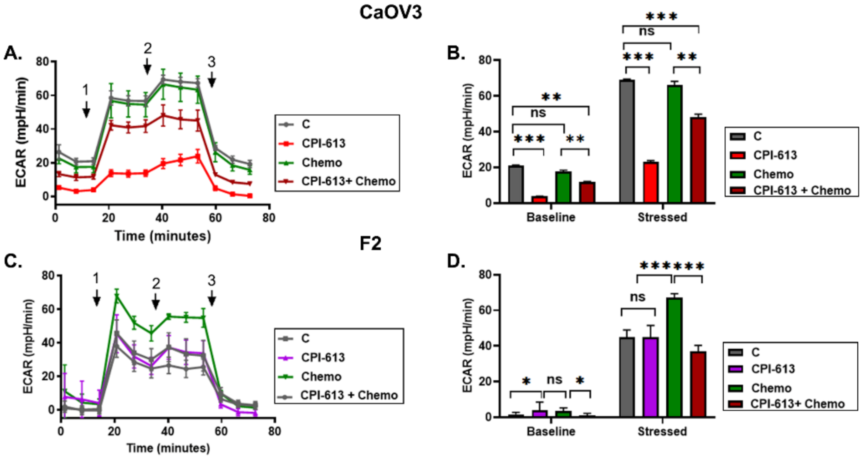

**Figure S4: CPI-613 inhibits mitochondrial energy and metabolism gene expression**

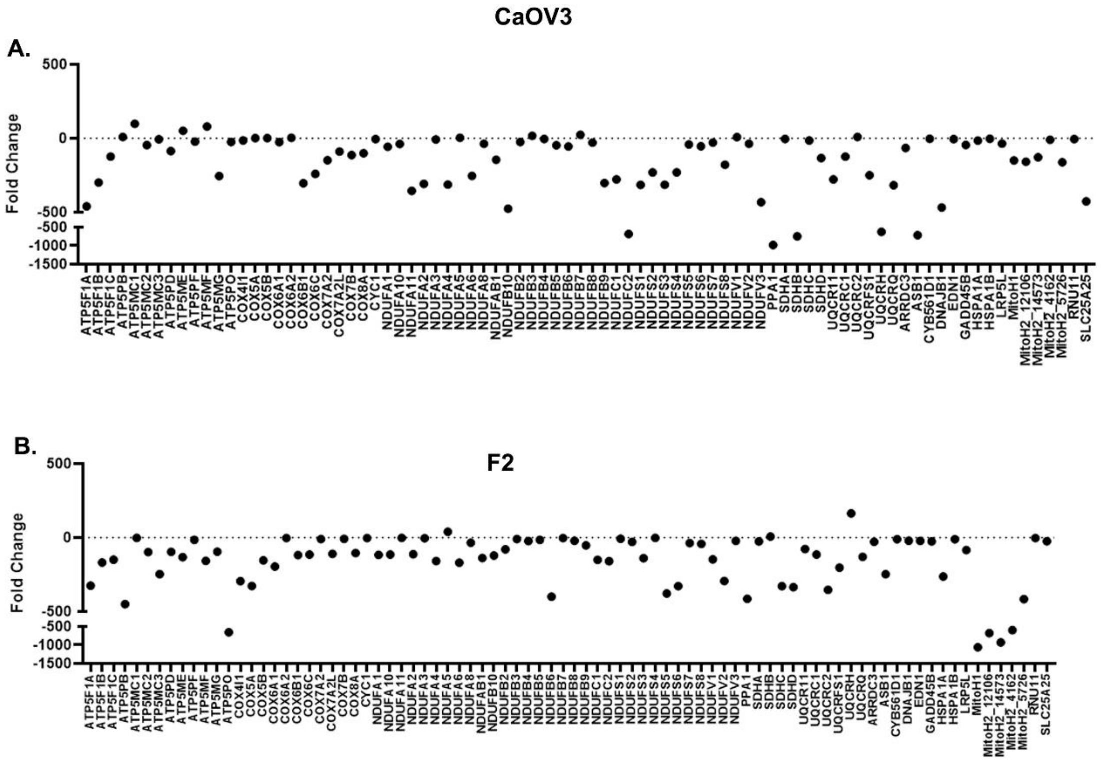

**Figure S5: CPI-613 modulates TCA metabolites in ovarian xenografts.**

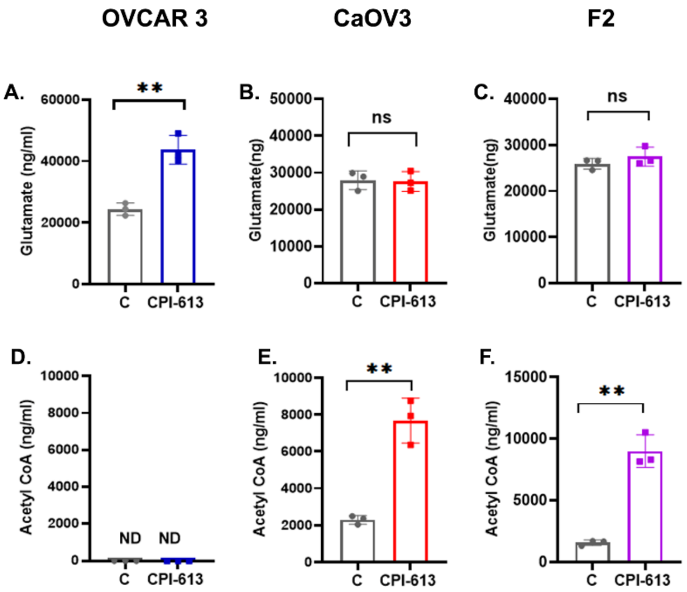
